# Priming anodal transcranial direct current stimulation attenuates short interval intracortical inhibition and increases time to task failure of a constant workload cycling exercise

**DOI:** 10.1101/2021.01.07.425802

**Authors:** Simranjit K Sidhu

## Abstract

Transcranial direct current stimulation (tDCS), a non-invasive neuromodulatory technique has been shown to increase the excitability of targeted brain area and influence endurance exercise performance. However, tDCS-mediated interaction between corticospinal excitability, GABA_A_ mediated intracortical inhibition and endurance exercise performance remains understudied. In two separate sessions, twelve subjects performed fatigue cycling exercise (80% peak power output) sustained to task failure in a double-blinded design, following either ten minutes of anodal tDCS (atDCS) or sham. Corticospinal excitability and short interval intracortical inhibition (SICI) were measured at baseline, post neuromodulation and post-exercise using paired-pulse transcranial magnetic stimulation (TMS) in a resting hand muscle. There was a greater a decrease in SICI (*P* < 0.05) post fatigue cycling with atDCS priming compared to sham. Time to task failure (TTF) was significantly increased following atDCS compared to sham (*P* < 0.05). These findings suggest that atDCS applied over the motor cortex can augment cycling exercise performance; and this outcome may be mediated via a decrease in the excitability of GABA_A_ inhibitory interneurons.

## INTRODUCTION

Central fatigue is manifested in a reduced ability of the central nervous system to fully activate a muscle or a muscle group during locomotor whole body exercise (Sidhu et al. 2014b). Recent studies have demonstrated that along with the development of central fatigue during acute locomotor exercise, the excitability of corticospinal neurons projecting to exercised lower limb and unexercised upper limb muscles is attenuated (Sidhu et al. 2014a; Weavil et al. 2016; Sidhu et al. 2017b). There is also some evidence to demonstrate that there is an increase in GABA_A_ (Sidhu et al. 2013) and GABA_B_ (Sidhu et al. 2018) mediated inhibition during cycling exercise sustained to task failure. This increase in intracortical inhibition may contribute to an attenuation in corticospinal excitability, leading to suboptimal endurance exercise performance (Sidhu et al. 2014a; Sidhu SK 2016).

Transcranial direct current stimulation (tDCS) is a non-invasive neuromodulation technique that delivers continuous low intensity electrical currents, causing significant changes in cortical excitability (Nitsche and Paulus 2000). Specifically, the application of tDCS is known to induce shifts in neuronal membrane excitability. For instance, *anodal tDCS* (*atDCS*) increases resting membrane potential, and *cathodal tDCS* (*ctDCS*) hyperpolarizes resting membrane potential (Nitsche et al. 2003a). Therefore, by mediating almost immediate changes in membrane excitability, tDCS impacts the response of involved neuronal circuits to incoming inputs. tDCS has the potential to attenuate the development of fatigue by moderating the way a given brain area processes a stimulus. For example, application of anodal tDCS on M1 for ten minutes prior to exercise increases time to task failure of a sustained isometric elbow flexion (Cogiamanian et al. 2007; Angius et al. 2016) and improve locomotor exercise performance (e.g. Vitor-Costa et al. 2015; Angius et al. 2018a; Lattari et al. 2018).

Short-interval intracortical inhibition (SICI), mediated via low-threshold GABA_A_ intracortical interneurons, can be measured with a form of paired-pulse TMS paradigm (Kujirai et al. 1993a). In this paradigm, a subthreshold conditioning stimulus is used to suppress the size of motor-evoked-potential (MEP) response produced by a suprathreshold stimulus given 1-5 ms later (Kujirai et al. 1993b). The application of atDCS for 10-20 min attenuates SICI measured in resting hand muscles (Cengiz et al. 2013). It is possible that the atDCS mediated effect on SICI may facilitate the increase in whole body exercise performance, however this remains unstudied.

The main objective of the current study was to examine the effect of atDCS on a) cycling exercise performance and b) SICI measured in an unexercised hand muscle from pre-post fatiguing cycling exercise sustained to task failure. We measured central responses in an unexercised remote muscle to examine the effect of fatiguing exercise on global cortical excitability. We hypothesised that while cycling exercise sustained to task failure will attenuate corticospinal excitability and increase SICI, the application of atDCS prior to exercise would attenuate the decrease in corticospinal excitability, ameliorate the increase in SICI, and increase the time to task failure (TTF) of a constant workload cycling exercise.

## METHODS

### Subjects

Twelve young (20.8 ± 0.4 years) healthy subjects were recruited for the study. All subjects were right handed (handedness laterality index: 0.86 ± 0.02) in accordance with the Edinburgh Handedness Inventory (Oldfield 1971). Participants with history of epilepsy, stroke, neurological illness, or those who were consuming psychoactive medications at the time of the experiments, were excluded from participation. Participants provided their written informed consent prior to participation and were instructed to avoid any strenuous activity at least 24 hrs prior to the experimental sessions. The study was approved by the University of Adelaide Human Research Ethics Committee and conducted in accordance with the Declaration of Helsinki.

### Experimental set-up and protocol

Participants were involved in 3 experimental sessions and each session was separated by at least five days to prevent any carry-over effects of neuromodulation and exercise-related fatigue. Each experimental session was performed between 11 am to 5 pm and, repeat sessions for each subject were conducted at the same time of the day to minimise the confounding influence of diurnal variations in cortisol on corticospinal excitability (Sale et al. 2008).

During the first session, participants were screened for any underlying health issues and familiarised with the experimental procedures. Participants then performed an incremental exercise test on a mechanically braked cycle ergometer (Velotron, Elite Model, Racer Mate, Seattle, WA) to establish peak power output (W_peak_). An initial cycling intensity of 25 W was increased by 25 Wmin^−1^ until task failure (Sidhu et al. 2014a). Task failure was defined as the inability to maintain 80% of the target cadence for more than 10 s despite verbal encouragement (Sidhu et al. 2014a). Subjects were instructed to maintain a cadence of 80 repeats per minute throughout the study.

In the followed-up 2 sessions (Fig. 1), subjects underwent either atDCS (10 minutes of atDCS at 2 mA) (Angius et al. 2016) or placebo (sham; 30 seconds stimulation at 2 mA to ensure the sensation associated with tDCS was apparent to the participant, followed by 9.5 min of no stimulation), in a counter-balanced, double-blind design. A set of stimulations consisting of 20 TMS (10 unconditioned and 10 conditioned responses randomly elicited to measure corticospinal excitability and SICI) and two motor nerve stimulations (MNS) were given at 5 time points: baseline, post neuromodulation (NM), 0, 10 and 20 min post fatigue cycling. During cycling exercise, heart rate (HR) (FT1, Polar Electro, Finland) and rating of perceived exertion (RPE) (modified Borg 0-10) were recorded every minute.

**Figure 1.**
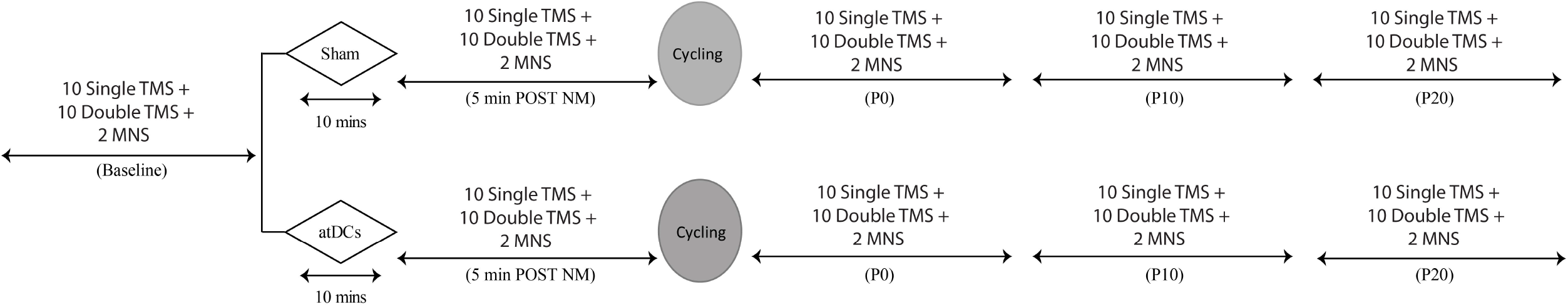
Schematic of experimental protocol. In sessions 2 and 3, subjects were given a dose of either atDCS (10 min @ 2mA) or sham (30 s stimulation + 9.5 min no stimulation) NM before performing the fatigue cycling exercise. A set of stimulations consisting of 10 single pulse TMS, 10-paired pulse TMS and 2 motor nerve stimulations (MNS) were applied at five time points (baseline, post neuromodulation (NM), 0 min post cycling exercise (P0), 10 min post cycling exercise (P10) and 20 min post cycling exercise (P20) in a resting FDI muscle at random order and each stimulation was separated by at least 4-5 seconds.

### Cycle ergometer setup

Subjects were set-up on a cycle ergometer with their feet strapped into the pedals while their right arm rested on a custom-made mould securely attached to the handle of the ergometer. Subjects were asked to restrict motion of the wrist and forearm, but the index finger was free to move. Subjects were also reminded to maintain a stable and constant upper body posture throughout the experimental session. This enabled a consistent and reliable application of TMS over the optimal location on the motor cortex.

### Electromyography recordings

After skin preparation with abrasion, alcohol swabs and shaving, surface electrodes were positioned over the muscle belly and tendon of the right first dorsal interosseous (FDI). EMG was recorded via monopolar configurations (Ag-AgCl, 10 mm diameter, inter-electrode distance: 2.9 ± 0.1 cm). EMG signals were amplified (100-1000 times; MA300 DTU, Motion Lab Systems, USA), band pass filtered (30-1000 Hz) and analogue to digitally converted at a sampling rate of 2000 Hz using a 16-bit power 1401 and Spike 2 data collection software (Cambridge Electronic Design, UK) via custom written scripts. Collected data was stored on a laboratory PC for offline analysis.

### Motor nerve stimulation

While the subjects were seated on the cycle ergometer, the optimal location at the ulnar nerve (proximal to the wrist joint) for eliciting muscle responses (M-waves) was established by applying low intensity single-pulse electrical stimuli (10 mA) using a constant voltage stimulator (model DS7AH, Digitimer Ltd., UK). The chosen stimulation location was one that produced the largest M-wave. The optimal stimulation intensity was determined by increasing the stimulus intensity by increments of 5 mA until the size of the M-wave demonstrated no further increase (i.e. producing maximal compound muscle activation potential; M_max_). The stimulation intensity was then increased by a further 20% to ensure maximal activation of the target muscle throughout the session (stimulation intensity: 18.3 ± 0.8 mA).

### Transcranial magnetic stimulation

Single and paired-pulse TMS were delivered to the left M1 using two MagStim 200 magnetic stimulators connected to a Bistim unit (MagStim, Dyfed, UK) and a figure-of-eight coil. The coil was placed tangentially to the scalp with the handle positioned approximately 45° posteriorly producing a current flow within brain in the posterior-anterior direction. The optimal location that gave the largest MEPs in the resting FDI muscle was marked directly on the scalp to ensure consistent positioning of the coil throughout the experiment as determined via visual inspection by the experimenter.

Active motor threshold (AMT) was measured as the minimum stimulus intensity required to elicit an MEP response in 5 out of 10 trials, while participants produced 10% of the their maximum voluntary contraction (MVC) force. Force feedback and target force (10% MVC) was provided on a monitor in front of the subject.

SICI was measured using a paired-pulse TMS paradigm consisting of a suprathreshold test stimulus (TS) preceded by a subthreshold conditioning stimulus with an interstimulus interval (ISI) of 3 ms (Kujirai et al. 1993b). The TS was set (81.54 ± 2.73) to produce an MEP with peak-to-peak amplitude of 1 mV in a resting FDI muscle (i.e. unconditioned MEP). The CS was set (39.58 ± 1.4) at an intensity (70%, 80% or 90% of AMT) which produced closest to 50% of unconditioned MEP (i.e. conditioned MEP) (Otieno et al. 2020).

### Anodal transcranial direct current stimulation

atDCS or sham were delivered through a direct current electrical stimulator (NeuroConn DC-Stimulator, Germany) connected to a pair of saline soaked electrodes (25 cm^2^). Anode was placed over the left motor cortex (hot spot determined by TMS) and cathode on the right supraorbital area (Waters-Metenier et al. 2014). Stimulation was applied at an intensity of 2 mA for a period of 10 min (atDCS) (Angius et al. 2016; Angius et al. 2018a) or 30 s followed by 9.5 min of no stimulation (sham). In order to minimise the discomfort caused by electrical transients, 10 s of fade in and fade out current was set at the start and end of the stimulation time (Nitsche et al. 2008). The selection of either atDCS or sham applied in the two experimental sessions was double blinded to both the subject and the main experimenter.

### Data analysis

Spike software (Cambridge Electronics Design, UK) was used for offline analysis of the EMG data. Since the aim was to quantify corticospinal excitability and SICI in a resting muscle, where muscle activity was more than 10 μV in the 100 ms prior to stimulation, data was removed from the analysis (< 3% trials) (Otieno et al. 2019). M_max_ and MEP amplitude was measured as peak-to-peak and expressed in mV. MEP was normalised to M_max_ at each time point to account for muscle-dependant changes (Todd et al. 2003). SICI was quantified by expressing conditioned MEP as a percentage of the average unconditioned MEP amplitude at each time point. Therefore, an increase in short-interval inhibition is demonstrated by a reduction in the magnitude of the measurement. Effects of NM on corticospinal excitability (i.e. MEP), SICI and M_max_ was investigated relative to the baseline. To assess the effects of fatigue, data obtained post fatigue cycling (i.e. P0, P10 and P20) was normalised to the average post NM for both sessions.

### Statistical analysis

One-way repeated measures analysis of variance (ANOVA) were used to compare RMT, AMT, M_max_, stimulation intensity required for a 1 mV MEP response (S1mV), baseline MEP, HR and RPE between sessions (i.e. sham; atDCS). Linear mixed model analyses with repeated measures were used to investigate the effect of time (i.e. baseline, post NM, P0, P10 and P20) and intervention (i.e. sham, atDCS) on MEP (% M_max_), SICI and M_max_. For linear mixed model analyses, subjects were included as random effect, and significant main effects and interactions were further investigated using custom contrasts with Bonferroni correction. Paired t-test was used to determine difference in time to task failure (TTF) between sessions. G * Power software was used to calculate Cohen’s effect sizes (dz) where statistical significance was detected with post hoc comparisons. Data is shown as mean ± standard error of the mean (SEM). Statistical significance is set at *P* < 0.05.

## RESULTS

All participants completed the study with no adverse reaction. There was no difference in baseline RMT, AMT, M_max_, S_1mv_ and SICI intensities between sessions (*F_1, 22_* < 0.4, *P* > 0.5; *Table 1*). In addition, no difference in baseline MEP amplitude at S_1mv_ was observed between sham and atDCS sessions (*F_1, 21_* = 0.13, *P* > 0.7; *Table 1*).

**Table 1.** Subject characteristics in Mean ± SEM at baseline

|  | Sham | atDCS |
| --- | --- | --- |
| RMT (% MSO) | 60 ± 1.6 | 59.7 ± 1.7 |
| AMT (% MSO) | 47.4 ± 1.3 | 47.6 ± 1.6 |
| SICI (conditioning, %AMT) | 38.6 ± 1.3 | 40.6 ± 1.5 |
| SICI (test) | 70.9 ± 2.9 | 83.8 ± 2.6 |
| S<sub>1mv</sub> | 79.1 ± 2.7 | 83.2 ± 2.6 |
| Mmax (mV) | 15.3 ± 0.9 | 13.4 ± 0.7 |
| Baseline MEP amplitude (mV) | 1.4 ± 0.08 | 1.3 ± 0.1 |

### Effect of atDCS

Figure 2 illustrates the changes in MEP (%M_max_) and SICI post NM for both sham and atDCS sessions. MEP changed with time (*F_1, 384_* = 8.2, *P* < 0.01) and NM (*F_1, 127_*= 13.5, *P* < 0.001). In addition, there was an interaction between time and NM (*F_1, 397_*= 10.9, *P* < 0.01). While MEP was attenuated after sham stimulation compared to baseline (*P* < 0.001), it did not change post atDCS from baseline (*P* = 0.74). In addition, MEP post NM was greater in the atDCS session compared to sham session (P < 0.001; Fig. 2A).

**Figure 2.**
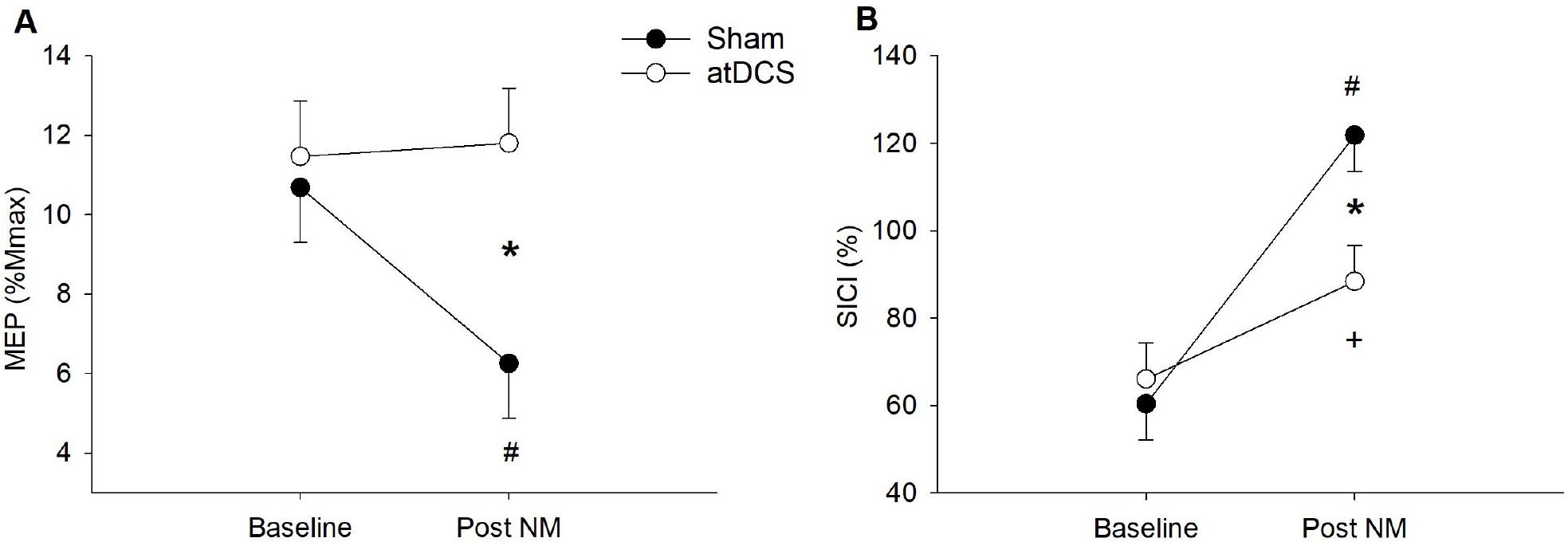
A: Corticospinal excitability (i.e. MEP % M_max_) and **B**: SICI post NM. *P<0.05 between sessions; #P<0.05 from baseline in sham and ^+^P<0.05 from baseline in atDCS session.

SICI changed with time (F_1,440_ = 57.9, P < 0.001) and NM (F_1,145_ = 5.4, P < 0.05) and an interaction between time and NM was also observed (F_1, 448_ = 12.5, P < 0.001). SICI was reduced after both sham (P < 0.001) and atDCS (P < 0.01). SICI was also less in the sham session compared to atDCS session (P < 0.001; Fig. 2B), suggesting an attenuation in the magnitude of inhibition with sham compared to atDCS.

M_max_ modulated with time (Baseline: 13.9 ± 1.2 mV; post NM: 13.5 ± 1.2 mV; *F_1, 63_= 5.7, P* < 0.05). However, no changes in M_max_ was seen with NM (*F_1, 12_* = 3.3, *P* = 0.10) and there was no interaction between time and NM (*F_1, 65_ = 2, P* = 0.17).

### Effect of Fatigue Cycling Exercise

Figure 3 A and B demonstrates corticospinal excitability (i.e. MEP %M_max_) and SICI respectively following fatigue cycling exercise measured at 0 min, 10 min and 20 min post exercise (i.e. P0, P10, P20) normalised to responses post NM in each experimental session. MEP (Fig. 3A) was not different across time (*F_2, 552_* = 1.5, *P* = 0.2) and there was no interaction between time and NM (*F_2, 529_* = 2.5, *P* = 0.1) However, there was a main effect of NM on MEP (average during sham: 115.3 ± 9.5 %; average during atDCS: 98.5 ± 9.5 %; *F_1, 260_* = 8.45, *P* < 0.01; Fig. 3A).

**Figure 3.**
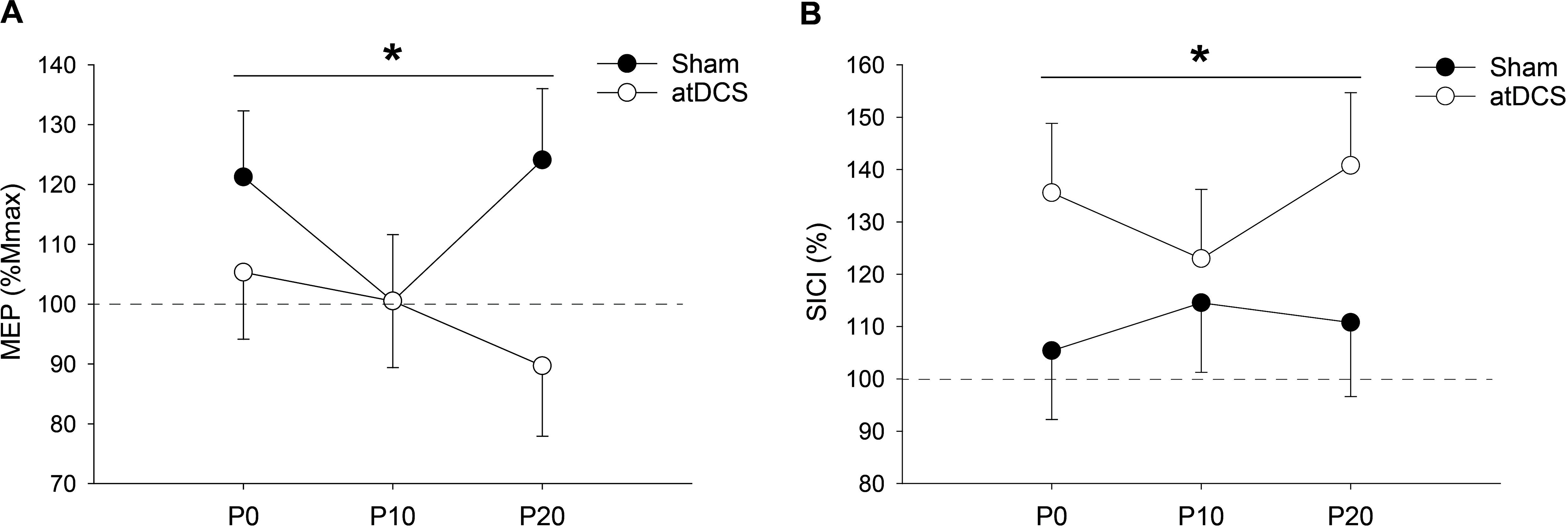
**A:** Changes in corticospinal excitability (i.e. MEP % M_max_) post FC normalised to average post NM. **B:** SICI post FC normalised to average post NM. *P<0.05 main effect of session.

SICI (Fig. 3B) did not change across time (*F_2, 516_* = 0.35, *P = 0.7*) and there was no interaction between time and NM (*F_2, 584_*= 0.98, *P = 0.37*). However, there was a main effect of NM on SICI (*F_1, 169_* = 6.8, *P* < 0.05; *Fig. 3B*) whereby the magnitude of inhibition was greater in sham compared to atDCS (Sham: 110.2 ± 11.4 %; atDCS: 133.1 ± 11.4 %)

M_max_ changed across time (PO: 95.1 ± 2.8%; P10: 92.4 ± 2.8%; P20: 89.7 ± 2.8%; *F_2, 91_* = 4.9, P < 0.01) and NM (Sham: 95.9 ± 3.1 %; atDCS: 88.9 ± 3.1 %; *F_1, 25_* = 6.8, *P* < 0.05) but no interaction was observed between time and NM (*F_2, 113_* = 1.27, *P* = 0.28).

### Effect of atDCS on TTF of Fatigue Cycling

TTF (min) improved after atDCS compared to sham (sham: 7.92 ± 0.5 min; atDCS: 9.71 ± 0.8 min; *P* <0.05, dZ = 4.52). The scatter plot in Figure 4 shows the % improvement in TTF after atDCS compared to sham whereby 11 out of 12 participants showed an improvement in TTF after atDCS compared to sham.

**Figure 4.**
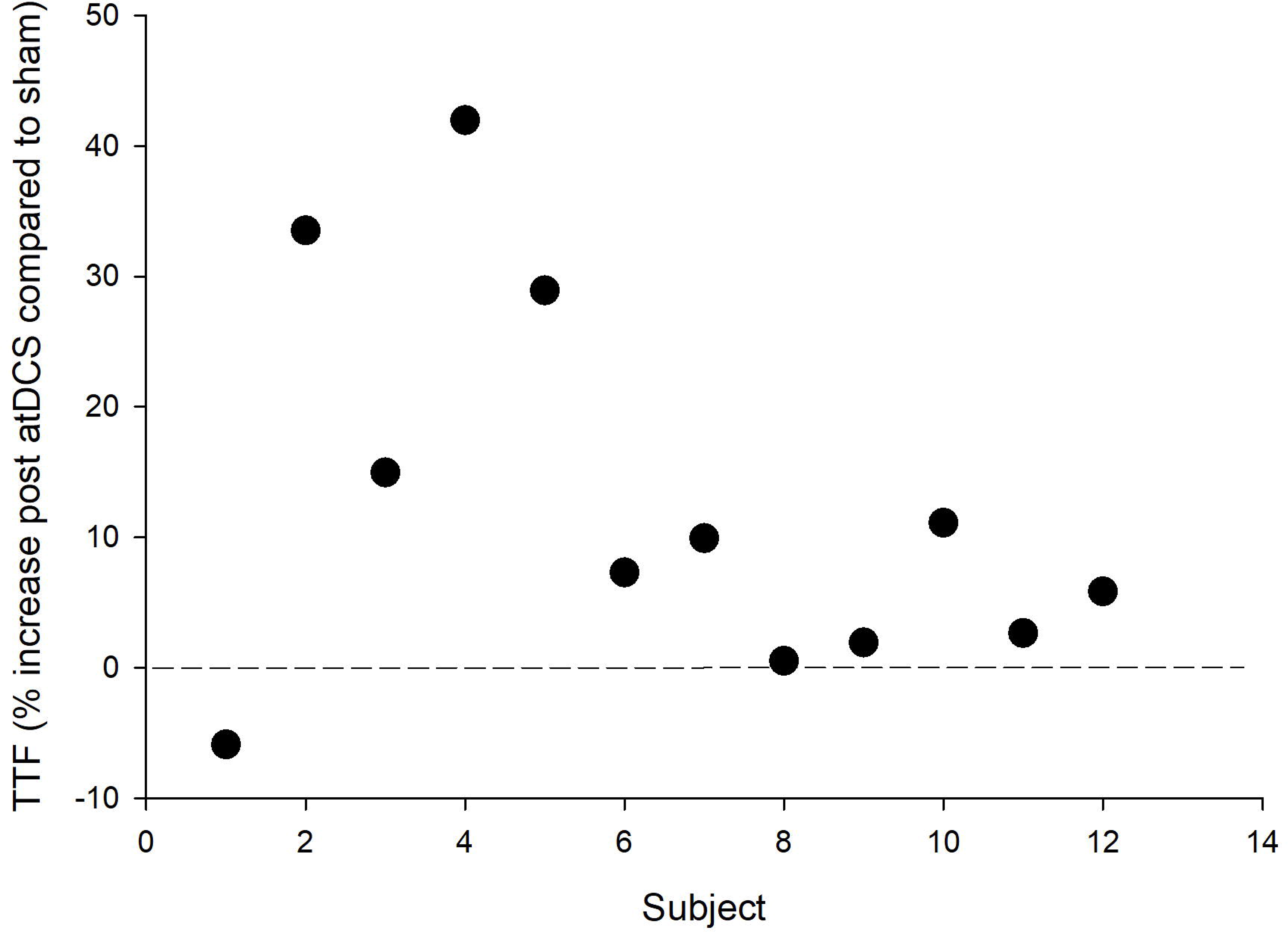
Individual TTF (% change) post atDCS session compared to the sham condition.

### HR and RPE

Table 2 shows the HR and RPE values for the first 5 minutes of fatigue cycling where every minute is an average of the values obtained from 12 subjects. First 5 minutes of the exercise is shown as all subjects managed to complete a minimum of five minutes of cycling. No significant differences was observed in HR and RPE between sham and atDCS sessions.

**Table 2.** Changes in group mean HR and RPE during cycling exercise (Mean ± SEM). Group mean data during first 5 minutes of exercise is shown as all subjects managed to complete at least 5 min of the exercise.

| Time (min) | HR (sham) | HR (atDCS) | RPE (sham) | RPE (atDCS) |
| --- | --- | --- | --- | --- |
| Rest | 69.0 ± 0.3 | 70.5 ± 0.4 | | |
| 1 | 145.8 ± 0.5 | 142.6 ± 0.6 | 4.2 ± 0.1 | 3.2 ± 0.1 |
| 2 | 158.9 ± 0.3 | 160.1 ± 0.3 | 6.3 ± 0.1 | 5.3 ± 0.1 |
| 3 | 168.4 ± 0.3 | 168.2 ± 0.3 | 7.6 ± 0.1 | 6.7 ± 0.1 |
| 4 | 174.3 ± 0.3 | 174.2 | 8.8 ± 0.0 | 8.1 ± 0.1 |
| 5 | 179.0 ± 0.2 | 179.3 ± 0.3 | 9.3 ± 0.0 | 8.9 ± 0.1 |
| . | . | . | . | . |
| . | . | . | . | . |
| . | . | . | . | . |
| Last Minute | 187 ± 0.3 | 187 ± 0.3 | 9.9 ± 0.0 | 10 ± 0.0 |

## DISCUSSION

### Main Findings

The current study aimed to investigate the effects of tDCS on motor cortical mechanisms including corticospinal tract excitability and SICI with fatigue cycling exercise, and the potential link between these underlying neural mechanisms and cycling exercise performance. The main outcome of the study was that atDCS priming attenuated the magnitude of SICI measured in a remote hand muscle post fatiguing cycling exercise, and augmented cycling exercise performance. The outcomes suggest that the atDCS mediated increase in whole body exercise performance may be mediated by cortical inhibitory mechanisms.

### Effects of priming atDCS on corticospinal excitability and SICI

tDCS involves the application of weak electric currents on the scalp that can be used to effectively shift transmembrane neuronal potentials via LTP (anodal) and LTD (cathodal) -like processes (Nitsche and Paulus 2001; Bailey et al. 2016), and influence corticospinal excitability (Nitsche et al. 2008). In the current study we found that atDCS applied for 10 minutes ameliorated the reduction in MEP seen with sham tDCS in a hand muscle. It is not clear why corticospinal excitability was attenuated after sham stimulation and tDCS did not produce LTP-like effect (i.e. an increase in corticospinal excitability demonstrated with an increase in MEP size). One factor underlying these observations may be the increased within and between sessions variability when using MEPs to measure changes in corticospinal excitability (Goldsworthy et al. 2016). The rather low number of stimulations delivered at each time point small may also have increased variability (Biabani et al. 2018). While we only had 12 participants in the current study, sample sizes ranging from 6 – 13 is common in tDCS studies involving exhaustive exercise (Okano et al. 2015; Abdelmoula et al. 2016; Angius et al. 2016; Barwood et al. 2016; Angius et al. 2018a; Baldari et al. 2018; Lattari et al. 2018). In addition, a range of neuromodulatory interventions have revealed a limited capacity to induce LTP-like modulation even 25 mins post priming (Muller-Dahlhaus et al. 2008; Sidhu et al. 2017a). The measurements in the current study were taken at 5 minutes post NM and it is possible that LTP-like plasticity effect may only consolidate over a longer period post NM.

SICI, a paired-pulse paradigm, provides a method to assess the responsiveness of the GABA_A_-ergic intracortical inhibitory cortical interneurons (Kujirai et al. 1993b; Hanajima et al. 1998). SICI decreases during voluntary contractions (Ridding et al. 1995; Soto et al. 2006) and this modulation is compatible with the increase in corticospinal excitability typically observed during voluntary contractions (Bawa and Lemon 1993; Di Lazzaro et al. 1998a). Therefore, it is conceivable that a decrease in MEP (reflective of the attenuated gain of the corticospinal pathway) represents an increase in the magnitude of SICI. In the current study, if anything, we observed a greater decrease in SICI with sham, compared to atDCS. The fact that there was greater SICI with atDCS compared to sham, even though the corresponding MEP remained unchanged, is indeed in contrary to what was expected (Nitsche et al. 2005). The underlying reason for this outcome is not clear, but could either be related to the time at which measurements are taken post tDCS or the large variability in response to tDCS protocols whereby 50% of individuals have been shown to have only a small or no response to tDCS as measured with TMS (Wiethoff et al. 2014).

### Effects of atDCS on exercise related RPE, HR, TTF, corticospinal excitability and SICI

The application of atDCS prior to exercise did not affect RPE and HR during the cycling exercise compared to the sham condition. This outcome is consistent with the findings of other studies that have also shown no change in maximal RPE and HR with cycling exercise after the application of tDCS (Okano et al. 2013; Vitor-Costa et al. 2015), even though one of these studies reported a slower rate of increase in RPE and a lower magnitude of HR during submaximal cycling workloads with atDCS (Okano et al. 2015). Our result demonstrated a significant increase in the TTF of the fatigue cycling exercise after the application of atDCS. This outcome is in agreement with other studies investigating the effects of atDCS on TTF of single-joint and whole body fatiguing exercises (e.g. Cogiamanian et al. 2007; Vitor-Costa et al. 2015; Abdelmoula et al. 2016; Angius et al. 2016; Angius et al. 2018b), but also counteracts other studies that have shown no effect of tDCS on TTF (Angius et al. 2015; Barwood et al. 2016; Baldari et al. 2018). The varying outcomes reported between studies may be related to a multitude of factors (Baldari et al. 2018); including differences in the tDCS montages used, for example monocephalic (Abdelmoula et al. 2016; Angius et al. 2018a) versus bicephalic montages (Okano et al. 2015; Vitor-Costa et al. 2015), single (Cogiamanian et al. 2007; Abdelmoula et al. 2016) versus double blinded (Williams et al. 2013; Angius et al. 2018a) study design, current magnitude and duration, the time during which atDCS is applied (before or during TTF) and the heterogeneity of participants including their fitness levels. The mechanisms of a-tDCS induced increase in exercise performance is not clear, however tDCS applied for 10-30 minutes has been shown to increase the excitability of the human motor cortex for at least an hour post stimulation (Nitsche and Paulus 2001). Therefore, it could be postulated that the atDCS mediated changes in intracortical mechanisms may play a role in influencing descending voluntary drive and exercise performance via changes in the development of muscular fatigue (Cogiamanian et al. 2007). Our results suggest that in addition to the fact that the priming with atDCS improved cycling exercise performance, it also attenuated the magnitude of SICI post exercise which remained lower compared to sham for at least 20 minutes post exercise. It is possible the atDCS mediated effects on the low threshold GABAa-ergic interneurons in the motor cortex during the cycling exercise resulted in an augmentation of performance. tDCS may have two possible effects on the cortical circuits that are tested with SICI. It can influence the corticospinal neurons (i.e. measured with MEP evoked by TMS) and/or it can influence the GABAa interneurons within the cortex. Our data show that priming with atDCS had no effect on MEP, even though SICI was reduced. These findings suggest that measures of corticospinal excitability and SICI are potentially mediated via independent mechanisms whereby an increase in SICI does not necessarily correspond to a decrease in the size of MEP and vice versa. In fact, it has been shown that even though cycling exercise modulates SICI, it does not necessarily influence MEP (Singh et al. 2014; Smith et al. 2014). Furthermore, when MEP is maintained constant (by adjusting the background contraction strength/EMG or stimulation intensity), modulation in intracortical inhibition is recorded (Sidhu et al. 2018). Indeed, an MEP is also influenced by spinal mechanisms (Taylor and Gandevia 2004), whereas SICI is known to be exclusively related to intracortical inhibitory mechanisms as demonstrated with pharmacological manipulations (Ziemann et al. 1996b; Ziemann et al. 1996a) and direct epidural recordings which show that the conditioning stimulus attenuated the magnitude of the late I-waves evoked by the test pulses (Di Lazzaro et al. 1998b). It is possible that tDCS had effects on spinal structures and or other cortical circuitry that influenced the MEP independently of the SICI mechanisms, although this remains a speculation and forms an important extension of work related to effects of tDCS on cortical versus spinal circuitry.

atDCS priming attenuated the MEP response measured after exercise in a remote muscle compared to the sham condition. The after-effects of atDCS appear to be subjected to the changes in the NMDA receptor efficacy similar to that demonstrated with pharmacological NMDA receptor modulation (Ziemann et al. 1998) rather than membrane polarisation which is effective during the stimulation (Nitsche et al. 2003b). The attenuation in MEP response post cycling exercise after the application of atDCS may also be related to previous history of activity within the targeted neuronal network. For example, neural potentiation is suppressed following high neuronal activity (e.g. priming with atDCS) but increased following low neuronal activity (i.e. sham in the current context) (Abraham 2008; Hulme et al. 2014), an outcome that represents homeostatic metaplasticity so as to maintain a balanced network function. Indeed, the relative reduction in corticospinal excitability and SICI in sham compared to atDCS post priming (Fig. 2) may partly explain the increase in corticospinal excitability and SICI post cycling exercise in the sham condition compared to atDCS.

In conclusion, this study provides evidence to demonstrate that atDCS applied prior to cycling exercise has the potential to improve constant load cycling exercise performance by increasing TTF and attenuate the magnitude of SICI post exercise for at least 20 minutes. These findings contribute to the current debate on the functional effectiveness of tDCS on motor performance and the neural circuitry during exhaustive cycling exercise.

## Funding

This research did not receive any specific grant from funding agencies in the public, commercial, or not-for-profit sectors

## Notes

### Competing Interest Statement

The authors have declared no competing interest.

## References

Abdelmoula A, Baudry S, Duchateau J (2016) Anodal transcranial direct current stimulation enhances time to task failure of a submaximal contraction of elbow flexors without changing corticospinal excitability. Neuroscience 322:94–103

Abraham WC (2008) Metaplasticity: tuning synapses and networks for plasticity. Nat Rev Neurosci 9:387–399 doi: 10.1038/nrn2356

Angius L, Hopker JG, Marcora SM, Mauger AR (2015) The effect of transcranial direct current stimulation of the motor cortex on exercise-induced pain. Eur J Appl Physiol 115:2311–2319 doi: 10.1007/s00421-015-3212-y

Angius L, Mauger AR, Hopker J, Pascual-Leone A, Santarnecchi E, Marcora SM (2018a) Bilateral extracephalic transcranial direct current stimulation improves endurance performance in healthy individuals. Brain Stimul 11:108–117 doi: 10.1016/j.brs.2017.09.017

Angius L, Mauger AR, Hopker J, Pascual-Leone A, Santarnecchi E, Marcora SM (2018b) Bilateral extracephalic transcranial direct current stimulation improves endurance performance in healthy individuals. Brain Stimulation: Basic, Translational, and Clinical Research in Neuromodulation 11:108–117

Angius L, Pageaux B, Hopker J, Marcora SM, Mauger AR (2016) Transcranial direct current stimulation improves isometric time to exhaustion of the knee extensors. Neuroscience 339:363–375

Bailey NW, Thomson RH, Hoy KE, Hernandez-Pavon JC, Fitzgerald PB (2016) TDCS increases cortical excitability: Direct evidence from TMS-EEG. Cortex 74:320–322 doi: 10.1016/j.cortex.2014.10.022

Baldari C, Buzzachera CF, Vitor-Costa M, Gabardo JM, Bernardes AG, Altimari LR, Guidetti L (2018) Effects of Transcranial Direct Current Stimulation on Psychophysiological Responses to Maximal Incremental Exercise Test in Recreational Endurance Runners. Front Psychol 9:1867 doi: 10.3389/fpsyg.2018.01867

Barwood MJ, Butterworth J, Goodall S, House JR, Laws R, Nowicky A, Corbett J (2016) The Effects of Direct Current Stimulation on Exercise Performance, Pacing and Perception in Temperate and Hot Environments. Brain Stimul 9:842–849 doi: 10.1016/j.brs.2016.07.006

Bawa P, Lemon RN (1993) Recruitment of motor units in response to transcranial magnetic stimulation in man. J Physiol 471:445–464

Biabani M, Farrell M, Zoghi M, Egan G, Jaberzadeh S (2018) The minimal number of TMS trials required for the reliable assessment of corticospinal excitability, short interval intracortical inhibition, and intracortical facilitation. Neurosci Lett 674:94–100 doi: 10.1016/j.neulet.2018.03.026

Cengiz B, Murase N, Rothwell JC (2013) Opposite effects of weak transcranial direct current stimulation on different phases of short interval intracortical inhibition (SICI). Exp Brain Res 225:321–331 doi: 10.1007/s00221-012-3369-0

Cogiamanian F, Marceglia S, Ardolino G, Barbieri S, Priori A (2007) Improved isometric force endurance after transcranial direct current stimulation over the human motor cortical areas. Eur J Neurosci 26:242–249 doi: 10.1111/j.1460-9568.2007.05633.x

Di Lazzaro V, Restuccia D, Oliviero A, et al. (1998a) Effects of voluntary contraction on descending volleys evoked by transcranial stimulation in conscious humans. J Physiol 508 (Pt 2):625–633

Di Lazzaro V, Restuccia D, Oliviero A, et al. (1998b) Magnetic transcranial stimulation at intensities below active motor threshold activates intracortical inhibitory circuits. Exp Brain Res 119:265–268 doi: 10.1007/s002210050341

Goldsworthy MR, Hordacre B, Ridding MC (2016) Minimum number of trials required for within- and between-session reliability of TMS measures of corticospinal excitability. Neuroscience 320:205–209 doi: 10.1016/j.neuroscience.2016.02.012

Hanajima R, Ugawa Y, Terao Y, Sakai K, Furubayashi T, Machii K, Kanazawa I (1998) Paired-pulse magnetic stimulation of the human motor cortex: differences among I waves. J Physiol 509 (Pt 2):607–618

Hulme SR, Jones OD, Raymond CR, Sah P, Abraham WC (2014) Mechanisms of heterosynaptic metaplasticity. Philos Trans R Soc Lond B Biol Sci 369:20130148 doi: 10.1098/rstb.2013.0148

Kujirai T, Caramia MD, Rothwell JC, et al. (1993a) Corticocortical inhibition in human motor cortex. J Physiol 471:501–519

Kujirai T, Caramia MD, Rothwell JC, et al. (1993b) Corticocortical inhibition in human motor cortex. J Physiol 471

Lattari E, de Oliveira BS, Oliveira BRR, de Mello Pedreiro RC, Machado S, Neto GAM (2018) Effects of transcranial direct current stimulation on time limit and ratings of perceived exertion in physically active women. Neurosci Lett 662:12–16 doi: 10.1016/j.neulet.2017.10.007

Muller-Dahlhaus JF, Orekhov Y, Liu Y, Ziemann U (2008) Interindividual variability and age-dependency of motor cortical plasticity induced by paired associative stimulation. Exp Brain Res 187:467–475 doi: 10.1007/s00221-008-1319-7

Nitsche MA, Cohen LG, Wassermann EM, et al. (2008) Transcranial direct current stimulation: State of the art 2008. Brain Stimul 1:206–223 doi: 10.1016/j.brs.2008.06.004

Nitsche MA, Liebetanz D, Antal A, Lang N, Tergau F, Paulus W (2003a) Modulation of cortical excitability by weak direct current stimulation--technical, safety and functional aspects. Suppl Clin Neurophysiol 56:255–276

Nitsche MA, Nitsche MS, Klein CC, Tergau F, Rothwell JC, Paulus W (2003b) Level of action of cathodal DC polarisation induced inhibition of the human motor cortex. Clin Neurophysiol 114:600–604 doi: 10.1016/s1388-2457(02)00412-1

Nitsche MA, Paulus W (2000) Excitability changes induced in the human motor cortex by weak transcranial direct current stimulation. J Physiol 527 Pt 3:633–639

Nitsche MA, Paulus W (2001) Sustained excitability elevations induced by transcranial DC motor cortex stimulation in humans. Neurology 57:1899–1901

Nitsche MA, Seeber A, Frommann K, et al. (2005) Modulating parameters of excitability during and after transcranial direct current stimulation of the human motor cortex. J Physiol 568:291–303 doi: 10.1113/jphysiol.2005.092429

Okano AH, Fontes EB, Montenegro RA, et al. (2015) Brain stimulation modulates the autonomic nervous system, rating of perceived exertion and performance during maximal exercise. Br J Sports Med 49:1213–1218 doi: 10.1136/bjsports-2012-091658

Okano AH, Fontes EB, Montenegro RA, et al. (2013) Brain stimulation modulates the autonomic nervous system, rating of perceived exertion and performance during maximal exercise. Br J Sports Med:bjsports-2012-091658

Oldfield RC (1971) The assessment and analysis of handedness: the Edinburgh inventory. Neuropsychologia 9:97–113

Otieno LA, Opie GM, Semmler JG, Ridding MC, Sidhu SK (2019) Intermittent single-joint fatiguing exercise reduces TMS-EEG measures of cortical inhibition. J Neurophysiol 121:471–479 doi: 10.1152/jn.00628.2018

Otieno LA, Semmler JG, Sidhu SK (2020) Single joint fatiguing exercise decreases long but not short-interval intracortical inhibition in older adults. Exp Brain Res doi: 10.1007/s00221-020-05958-w

Ridding MC, Taylor JL, Rothwell JC (1995) The effect of voluntary contraction on cortico-cortical inhibition in human motor cortex. J Physiol 487 (Pt 2):541–548 doi: 10.1113/jphysiol.1995.sp020898

Sale MV, Ridding MC, Nordstrom MA (2008) Cortisol inhibits neuroplasticity induction in human motor cortex. J Neurosci 28:8285–8293 doi: 10.1523/JNEUROSCI.1963-08.2008

Sidhu SK, Lauber B, Cresswell AG, Carroll TJ (2013) Sustained cycling exercise increases intracortical inhibition. Med Sci Sports Exerc 45:654–662 doi: 10.1249/MSS.0b013e31827b119c

Sidhu SK, Pourmajidian M, Opie GM, Semmler JG (2017a) Increasing motor cortex plasticity with spaced paired associative stimulation at different intervals in older adults. Eur J Neurosci 46:2674–2683 doi: 10.1111/ejn.13729

Sidhu SK, Weavil JC, Mangum TS, Jessop JE, Richardson RS, Morgan DE, Amann M (2017b) Group III/IV locomotor muscle afferents alter motor cortical and corticospinal excitability and promote central fatigue during cycling exercise. Clin Neurophysiol 128:44–55 doi: 10.1016/j.clinph.2016.10.008

Sidhu SK, Weavil JC, Thurston TS, et al. (2018) Fatigue-related group III/IV muscle afferent feedback facilitates intracortical inhibition during locomotor exercise. J Physiol 596:4789–4801 doi: 10.1113/JP276460

Sidhu SK, Weavil JC, Venturelli M, et al. (2014a) Spinal mu-opioid receptor-sensitive lower limb muscle afferents determine corticospinal responsiveness and promote central fatigue in upper limb muscle. J Physiol 592:5011–5024 doi: 10.1113/jphysiol.2014.275438

Sidhu SK, Weavil JC, Venturelli M, et al. (2014b) Spinal mu-opioid receptor-sensitive lower limb muscle afferents determine corticospinal responsiveness and promote central fatigue in upper limb muscle. J Physiol 592:5011–5024 doi: 10.1113/jphysiol.2014.275438

Sidhu SK WJ, Mangum TS, Jessop JE, Richardson RS, Morgan DE & Amann M (2016) Group III/IV locomotor muscle afferents alter motor cortical and corticospinal excitability and promote central fatigue during cycling exercise. Clin Neurophysiol doi: http://dx.doi.org/10.1016/j.clinph.2016.10.008

Singh AM, Duncan RE, Neva JL, Staines WR (2014) Aerobic exercise modulates intracortical inhibition and facilitation in a nonexercised upper limb muscle. BMC Sports Sci Med Rehabil 6:23 doi: 10.1186/2052-1847-6-23

Smith AE, Goldsworthy MR, Garside T, Wood FM, Ridding MC (2014) The influence of a single bout of aerobic exercise on short-interval intracortical excitability. Exp Brain Res 232:1875–1882 doi: 10.1007/s00221-014-3879-z

Soto O, Valls-Sole J, Shanahan P, Rothwell J (2006) Reduction of intracortical inhibition in soleus muscle during postural activity. J Neurophysiol 96:1711–1717 doi: 10.1152/jn.00133.2006

Taylor JL, Gandevia SC (2004) Noninvasive stimulation of the human corticospinal tract. J Appl Physiol 96:1496–1503

Todd G, Taylor JL, Gandevia SC (2003) Measurement of voluntary activation of fresh and fatigued human muscles using transcranial magnetic stimulation. The Journal of Physiology 551:661–671 doi: 10.1113/jphysiol.2003.044099

Vitor-Costa M, Okuno NM, Bortolotti H, Bertollo M, Boggio PS, Fregni F, Altimari LR (2015) Improving cycling performance: transcranial direct current stimulation increases time to exhaustion in cycling. PloS one 10:e0144916

Waters-Metenier S, Husain M, Wiestler T, Diedrichsen J (2014) Bihemispheric transcranial direct current stimulation enhances effector-independent representations of motor synergy and sequence learning. J Neurosci 34:1037–1050 doi: 10.1523/JNEUROSCI.2282-13.2014

Weavil JC, Sidhu SK, Mangum TS, Richardson RS, Amann M (2016) Fatigue diminishes motoneuronal excitability during cycling exercise. J Neurophysiol 116:1743–1751 doi: 10.1152/jn.00300.2016

Wiethoff S, Hamada M, Rothwell JC (2014) Variability in response to transcranial direct current stimulation of the motor cortex. Brain Stimul 7:468–475 doi: 10.1016/j.brs.2014.02.003

Williams PS, Hoffman RL, Clark BC (2013) Preliminary evidence that anodal transcranial direct current stimulation enhances time to task failure of a sustained submaximal contraction. PLoS One 8:e81418 doi: 10.1371/journal.pone.0081418

Ziemann U, Chen R, Cohen LG, Hallett M (1998) Dextromethorphan decreases the excitability of the human motor cortex. Neurology 51:1320–1324 doi: 10.1212/wnl.51.5.1320

Ziemann U, Lonnecker S, Steinhoff BJ, Paulus W (1996a) The effect of lorazepam on the motor cortical excitability in man. Exp Brain Res 109:127–135 doi: 10.1007/BF00228633

Ziemann U, Lonnecker S, Steinhoff BJ, Paulus W (1996b) Effects of antiepileptic drugs on motor cortex excitability in humans: a transcranial magnetic stimulation study. Ann Neurol 40:367–378 doi: 10.1002/ana.410400306

